# PMI estimation on aqueous humour: human translation from a metabolomic animal model

**DOI:** 10.1101/2025.04.15.648907

**Authors:** Alberto Chighine, Matteo Stocchero, Fabio De-Giorgio, Matteo Nioi, Ernesto d’Aloja, Emanuela Locci

**Affiliations:** Department of Medical Sciences and Public Health, Section of Legal Medicine, University of Cagliari, Cagliari, Italy; Department of Women’s and Children’s Health, University of Padova, Padova, Italy; Department of Health Surveillance and Bioethics, Section of Legal Medicine, Catholic University of Rome, Rome, Italy; Fondazione Policlinico Universitario A. Gemelli IRCCS, Rome, Italy

**Keywords:** Aqueous Humour, ^1^H NMR metabolomics, PMI, human translation, animal model

## Abstract

Translating findings from animal models to human applications remains a fundamental challenge across scientific research, with unique implications for post-mortem metabolomics. This study advances this goal by applying ^1^H NMR metabolomics to human aqueous humour, building on to a previously published sheep model for post-mortem interval estimation. We constructed a human dataset of 21 aqueous humor samples from 11 forensic autopsies, with post-mortem intervals (225– 1164 minutes) aligned with the animal model’s time range. Quantification of 46 metabolites revealed 43 shared between species, demonstrating qualitative similarities despite quantitative differences driven by species-specific factors, such as lactate and glutamate. Partial least squares regression models, which resulted highly accurate in the sheep model, showed increased prediction error in humans, underscoring translational complexities. Notably, taurine and hypoxanthine were identified as post-mortem interval-specific metabolites and, most of all, resulted unaffected by species, suggesting their relevance in the post-mortem interval maintained across species. This study is the first to attempt a translation of animal-derived ^1^H NMR metabolomic results to real-life human samples, addressing prior limitations through aligned timeframes and rigorous methodology. Animal models in post-mortem metabolomics offer controlled, reproducible data by minimizing real-life, providing a robust foundation for studying metabolomic modifications. However, direct translation to humans appears possible for a limited part of the metabolome, with key metabolites like taurine and hypoxanthine showing consistency. These findings endorse animal-derived metabolomics as a guide for human studies across diverse metabolomics investigations, promoting human studies on larger cohorts and more specific experimental designs.

## 1. Introduction

Despite considerable progress in forensic science, estimating the time since death, or post-mortem interval (PMI), remains a complex challenge. Accurate PMI determination is vital for forensic investigations, offering crucial insights for reconstructing event sequences and elucidating the conditions of death. However, the biological processes that occur during and after death are inherently complex, influenced by both environmental factors and individual differences, which significantly complicates PMI assessments [1].

The current gold standard for PMI estimation is Henssge’s nomogram, which evaluates post-mortem body temperature decline as it approaches equilibrium with the surrounding environmental temperature [2]. Other traditional methods such as the evaluation of *rigor* and *livor mortis* relies on subjective grading of post-mortem changes or empirical observations, lacking statistical validation, which affects their precision, reliability, and, ultimately, their use in court [3].

For these reasons, PMI estimation is one of the most investigated issues by the forensic researcher’s community with multiple applications of novel technologies and approaches [4].

Given that post-mortem tissues and biofluids undergo biological processes like autolysis and putrefaction, which progressively degrade their structure over time, a significant limitation is the choice of the biological matrix for analysis. This has led to the selection of more stable matrices, such as the ocular components [5], vitreous humor (VH) being traditionally utilized for forensic purposes and especially for PMI estimation [6]. The most fortunate attempt to estimate PMI on VH is based on the measurement of post-mortem potassium rise which has been demonstrated to present a linear increase after death, making it suitable for PMI estimation although its practical application is currently limited [7].

Among the novel approaches, metabolomics, based on the systematic study of low molecular weight metabolites, has been used to investigate PMI [8, 9]. The scientific rationale of post-mortem metabolomics in PMI investigation is that metabolome modifications occurring after death are mostly driven by PMI itself [10].

Our research group was one of the first to investigate PMI on ocular biofluids through a ^1^H Nuclear Magnetic Resonance (NMR) metabolomic approach on animal models [11-13]. In the hypothesis that immediate post-mortem transformation predominantly affects the anterior segment, where a very slight barrier, namely the cornea, separates the inner eye from the external environment, we initially examined the aqueous humour (AH), which is situated behind the cornea. AH, being acellulated, does not possess an inner metabolism, however, its metabolite content is well known in both animal and human setting [14, 15].

We studied post-mortem AH metabolomics on a highly controlled ovine model on a PMI window ranging from 118 to 1429 minutes [11]. In this experiment, we focused on two major variables, namely PMI and the effect of open and closed eyelids, demonstrating that the latter was negligeable with respect to PMI. A regression model for PMI estimation was built and validated with an independent set of samples resulting in an error in prediction of 99 minutes over the entire PMI window. While the approach was based on a profile of 43 quantified metabolites, a few of them (namely taurine, choline, and succinate) were found to be significantly correlated with PMI although individual models based on single metabolites resulted less performant than the entire profile in predicting PMI.

The same dataset was then challenged with the scientific gold standard for PMI estimation in ocular biofluids, i.e. potassium concentration, which turned out to be less efficient than metabolomics in predicting PMI across the same time frame [12]. Notably, when the PMI was divided into three different ranges, the prediction performance of metabolomics and potassium models diverged. The error associated with the metabolomic approach tended to increase over time, while the potassium method showed improvement, with both methods yielding superposable results for PMIs exceeding 1000 minutes.

The need for an animal model was due to two major experimental reasons: little inter-individual variability and the necessity of working on samples with known PMI. The experiment was indeed designed in order to maximise the focus on PMI related modifications in the AH metabolome and such conditions would have been realistically impossible to achieve working on human samples for several reasons related to practical and ethical issues. Indeed, sample size, AH withdrawal feasibility, known PMI, homogeneity in cause of death, sex, age, and pathological conditions – among all – are all limiting factors in designing post-mortem human research.

Furthermore, the evident inter-species differences and the enormous inter-individual variability characterizing humans must be taken into account. While the immediate human translation does not appear realistically achievable, investigating inter-species metabolomic behaviour could help in understanding post-mortem related modifications that may be ultimately used in bridging the gap between highly controlled experimental animal model and real-life setting.

In this paper, ^1^H NMR metabolomics was performed on AH human samples collected from real forensic cases with the aim of investigating post-mortem related metabolome modifications and comparing with those previously found in the animal model [11]. To the best of our knowledge, this represents the first post-mortem metabolomics experiment aimed at translating evidence gathered from an animal model to human samples.

## 2. Materials and methods

### 2.1 Samples collection and preparation

AH samples were collected during 11 consecutive judicial autopsies in medical malpractice cases performed at Forensic and Legal Medicine Institute of Catholic University of Rome. Approximately 0.5 ml of AH was withdrawn from eyes using a 5 ml syringe G22 through a corneal puncture in the anterior chamber passing through the limbus, at different post-mortem times, ranging from 225 to 1164 min (approximately 3.75–19.4 h).

Collection was not performed in case of pathological eye conditions and/or macroscopic blood contamination. After collection, AH samples were immediately stored at -80°C. Then, samples were shipped on dry ice to the Forensic and Legal Medicine Institute of University of Cagliari for the metabolomic analysis. Sampling was limited to the first 24 hours after death to match the PMI window of our previously investigated ovine AH model [11].

### 2.2 Sample preparation for NMR analysis

AH human samples were treated as previously described for the animal experiment [11]. Briefly, before the NMR analysis, samples were thawed and ultrafiltered with 10 kDa filter units (Amicon-10 kDa; Merck Millipore, Darmstadt, Germany) to remove proteins. 500 μl of filtered AH samples were then dried overnight using a Speed Vacuum concentrator (Eppendorf concentrator plus, Eppendorf AG, Hamburg, Germany), and reconstituted with 700 μl a 0.05 M phosphate buffer solution (pH 7.4) in D_2_O (99,9%, Cambridge Isotope Laboratories Inc, Andover, USA) containing the internal standard sodium 3-(trimethylsilyl) propionate-2,2,3,3,-d4 (TSP, 98 atom % D, Sigma-Aldrich, Milan) at a 0.2 mM final concentration.

### 2.3 ^1^H NMR experiments and data processing

^1^H NMR experiments were carried out on a Varian UNITY INOVA 500 spectrometer (Agilent Technologies, CA, USA) operating at 499.839 MHz, using the experimental conditions previously reported for ovine AH.

Spectra were processed using MestReNova software (Version 9.0, Mestrelab Research S.L.). After applying a 0.5 Hz line broadening of and 128K zero-filling the free induction decays (FID) were Fourier transformed. All spectra were phased, baseline corrected and referenced to TSP at 0.00 ppm. 46 metabolites were quantified by using the Chenomx NMR Suite Profiler tool (Chenomx 8.2 Inc., Edmonton, Canada – see Supplementary Table 1). The final dataset was submitted to for multivariate statistical analysis after Mean centring and unit variance scaling.

**Table 1.**
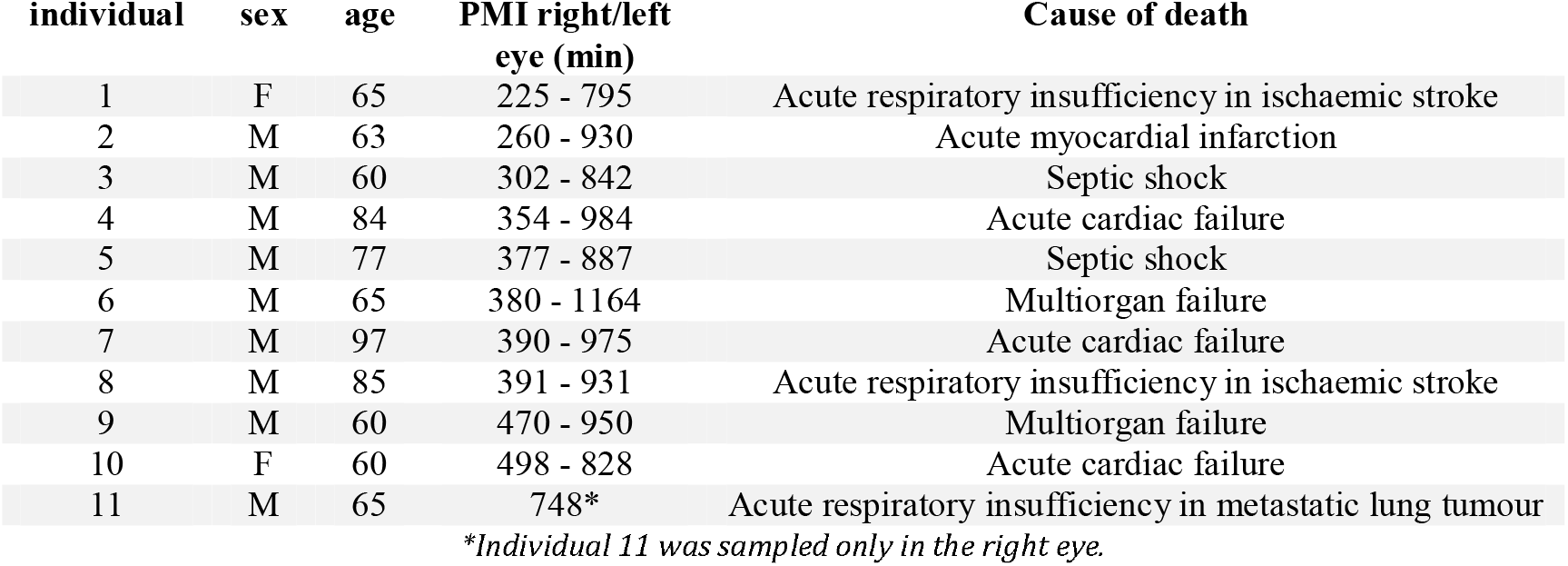
Demographic parameters, PMI, and causes of death.

| individual | sex | age | PMI right/left<br>eye (min) | Cause of death |
| --- | --- | --- | --- | --- |
| 1 | F | 65 | 225 - 795 | Acute respiratory insufficiency in ischaemic stroke |
| 2 | M | 63 | 260 - 930 | Acute myocardial infarction |
| 3 | M | 60 | 302 - 842 | Septic shock |
| 4 | M | 84 | 354 - 984 | Acute cardiac failure |
| 5 | M | 77 | 377 - 887 | Septic shock |
| 6 | M | 65 | 380 - 1164 | Multiorgan failure |
| 7 | M | 97 | 390 - 975 | Acute cardiac failure |
| 8 | M | 85 | 391 - 931 | Acute respiratory insufficiency in ischaemic stroke |
| 9 | M | 60 | 470 - 950 | Multiorgan failure |
| 10 | F | 60 | 498 - 828 | Acute cardiac failure |
| 11 | M | 65 | 748* | Acute respiratory insufficiency in metastatic lung tumour |

*\*Individual 11 was sampled only in the right eye.*

### 2.3 Statistical data analysis

Exploratory data analysis was performed by Principal Component Analysis (PCA) [16]. PLS regression [17] was applied to investigate the relationships between the metabolic content of the AH samples and PMI. Specifically, the number of score components to use was determined on the basis of the first maximum of the R^2^ calculated by 10-repeated 5-fold full cross-validation, i.e. Q^2^, under the constraint to pass the permutation test on the response. Stability selection based on variable selection considering the variable influence on projection parameter [18] was applied to discover the most relevant metabolites. A significance level of α=0.05 was assumed. PLS for designed experiments considering the design factor species, the design factor PMI and their interaction was applied to discover the effects of the design factors on the AH metabolome. An orthogonal design matrix was assumed. Model complexity was determined investigating the significance of the PLS-eigenvalues. The role played by each metabolite was evaluated calculating the Selectivity Ratio (SR) parameter. A significance level α=0.05 was assumed. The effects of species and PMI on the concentration of each single metabolite were investigated also by Multiple Linear Regression controlling the False Discovery Rate at level 0.10 by the Benjamini-Hochberg procedure [19]. To obtain a dataset balanced with respect to the factor species and with the same range of PMI for both animal and human samples, a procedure of sample selection was applied to the set of the animal samples. Specifically, the scores of the PLS regression model obtained considering only the AH animal samples within the range of PMI of the human samples and those of the PCA model of the residuals of that model were used to represent the animal samples. Then, hierarchical cluster analysis based on Ward’s method was applied to determine a number of clusters equal to the number of human samples and the animal sample closest to the center of each cluster was selected. Data analysis was performed by R-functions developed in-house using the R 4.2.2 platform (R Foundation for Statistical Computing).

## 3. Results

A total of 21 AH samples were collected from corpses of both gender (with a M to F ratio of approximately 1:5), aged from 60 to 97 years (mean=65, SD=14) with a PMI ranging from 225 to 1164 minutes (mean=684, SD=291) corresponding to 3.75 and 19.4 hours, respectively. Table 1 reports the demographic data, PMI, and cause of death.

The present experiment was consistently conducted using the same protocol applied to the animal dataset which consisted of 59 AH samples collected from young (24-48 months) ovis aries females, with PMI ranging from 118 to 1429, corresponding to 2 and 23.8 hours, respectively. ^1^H NMR spectra of animal and human AH showed very similar metabolomic composition with differences relying only in meabolite concentrations. A total of 46 endogenous metabolites were identified and quantified in human samples, of which 43 were shared with the animal samples. In particular, 2-Hydroxyisovalerate, Ethanolamine, and Glucose were identified only in human AH samples (Glucose was only present in few animal samples at PMI < 180 min). For the sake of comparison, the same 43 metabolites were considered in the following analyses.

An exploratory PCA analysis of both animal and human samples indicated in the score plot two clearly different clusters depending on the species under study (Figure 1A). Considering that the metabolites pool was the same, differences in the metabolomic profiles are hence only quantitative and mainly driven by Lactate, Glutamate, Glycine, Isoleucine and Leucine as can be see in PCA loading plot (Figure 1B).

**Figure 1.**
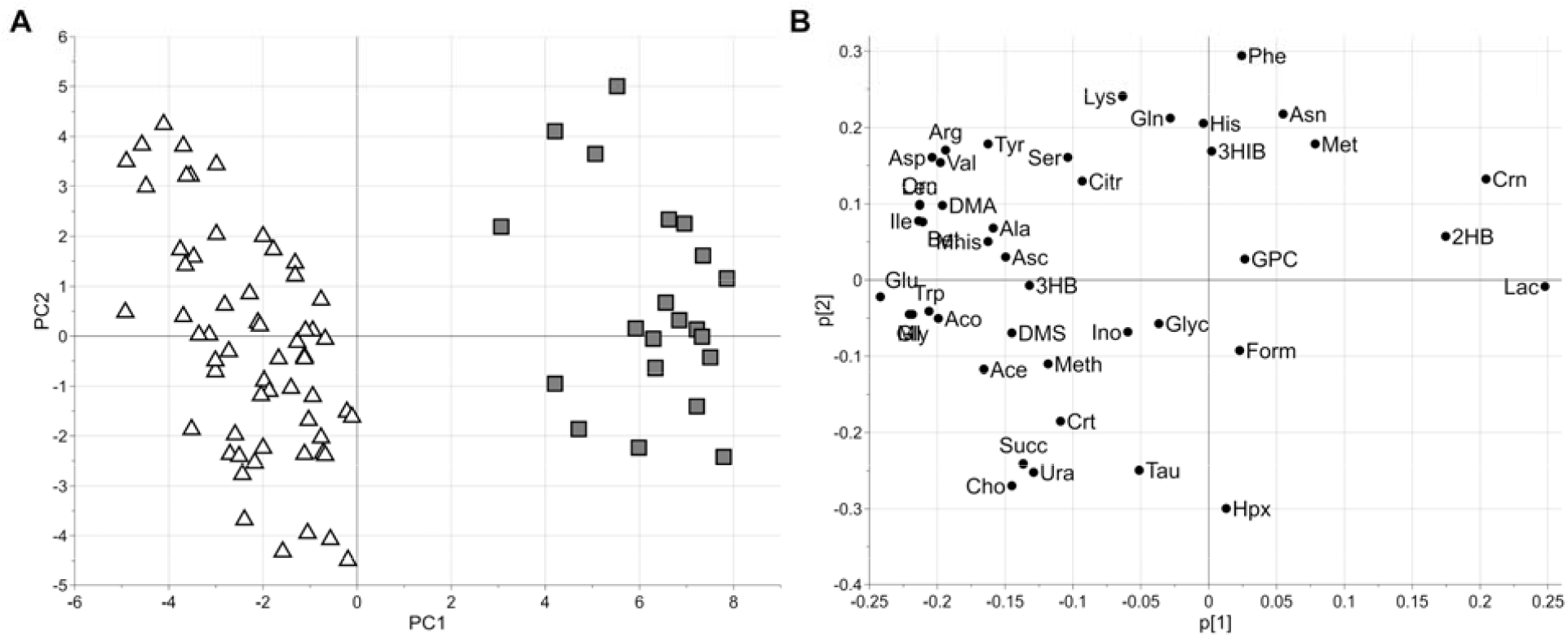
PCA model: score scatter plot obtained considering the first (PC1 explaining the 36% of the total variance) and the second (PC2 explaining the 11% of the total variance) score components (panel A) and the related loading plot (panel B). Triangles and boxes are used for Animal (A) and Human (H) AH samples, respectively.

The AH animal data were used to train a regression model for PMI estimation. The PLS model showed 2 score components, R^2^=0.949 (p=0.002), Q^2^=0.915 (p=0.002), standard deviation error in calculation (SDEC) equal to 87 minutes and standard deviation error in cross-validation (SDECV) equal to 112 minutes. The model was applied to predict the PMI of the AH human samples (Figure 2). Notably, human AHs display a larger data scattering indicative of the inherent variability in these samples. The standard deviation of the error in prediction (SDEP) for human samples resulted to be 300 minutes across the entire PMI window, highlighting that translating controlled animal experiments into the complexities of real-world human forensic scenarios is not directly achievable, despite supposed biological similarities. Of note, the SDEP resulted significantly higher than the SDECV and higher than the error estimated in prediction considering our previously published model, namely 99 minutes [11]. This comparison underscores the model’s robustness in controlled settings pointing out the need for further refinement to account for the diverse factors influencing PMI in human cases.

**Figure 2.**
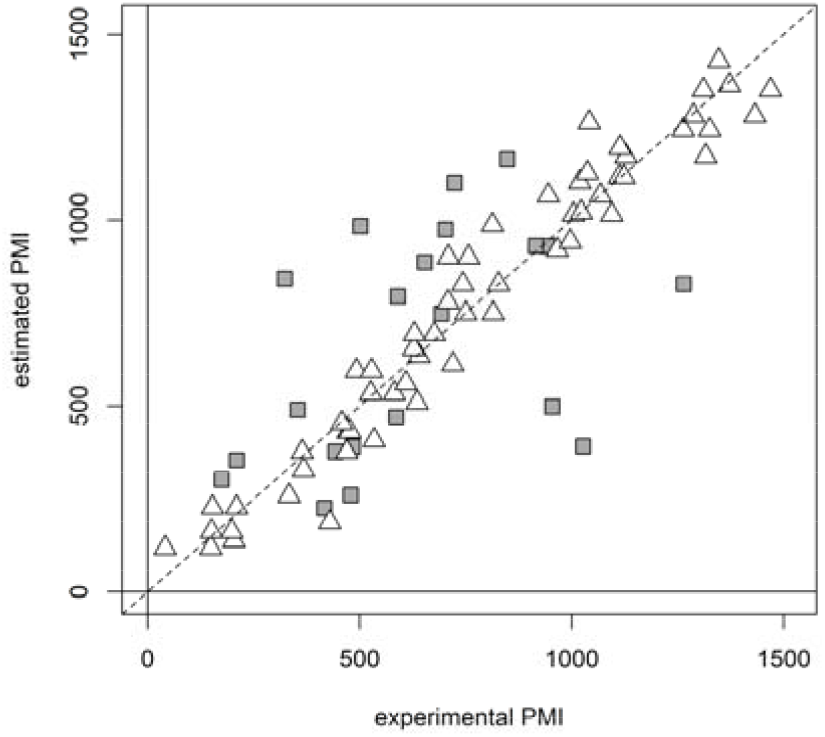
Regression model to estimate PMI based on the AH metabolomic data of the animal samples. PMI predicted for the AH human samples are reported as dark grey boxes whereas PMI calculated for the animal samples as white triangles. The dashed line represents the line of perfect agreement between estimated PMI and experimental PMI.

In order to better understand the prediction performance disparity, due to the difference between the animal and human datasets, we proceeded with animal sample selection to match the datasets for sample size and PMI window, aiming for a more direct comparison. This led to the selection of 21 out of 59 samples from the original animal model with a PMI range of 227 to 1127 minutes that were matched with the 21 human samples with a PMI range of 225 to 1164 minutes. The design matrix including the factors species and PMI and their interaction was approximately orthogonal.

A PLS for designed experiment model (Figure 3) was built to prove the relationships between PMI and species and AH metabolome, and to discover metabolomics features related to the two experimental factors. The model showed 1 score component explaining the factor PMI (R^2^=0.848, p<0.001), 1 score component for the factor species (MCC=1.000, p<0.001) whereas the interaction factor was not significant (p=0.133). Figure 3A displays a clear cluster structure associated to the species and trends depending on the PMI. Interestingly, while animal samples appear to maintain a linear behaviour over time, human samples tend to diverge at higher PMIs, reasonably due to the higher variability inside the human samples. Figure 3B, that shows the selectivity ratios (SR) related to the two investigated design factors, gives information about most significant metabolites that are associated with species differences or changes over time due to PMI. This differentiation is crucial for understanding how metabolomic profiles can be used to estimate PMI or identify species-specific markers in forensic science. Despite some metabolites like Lactate, Glutamate, and Isoleucine appear to be species-related being then crucial in distinguishing between human and animal samples, accordingly to the preliminary data analysis based on PCA, others (Choline and Taurine) appear to be intriguingly indifferent to the species and related exclusively to the metabolic changes occurring in the post-mortem.

**Figure 3.**
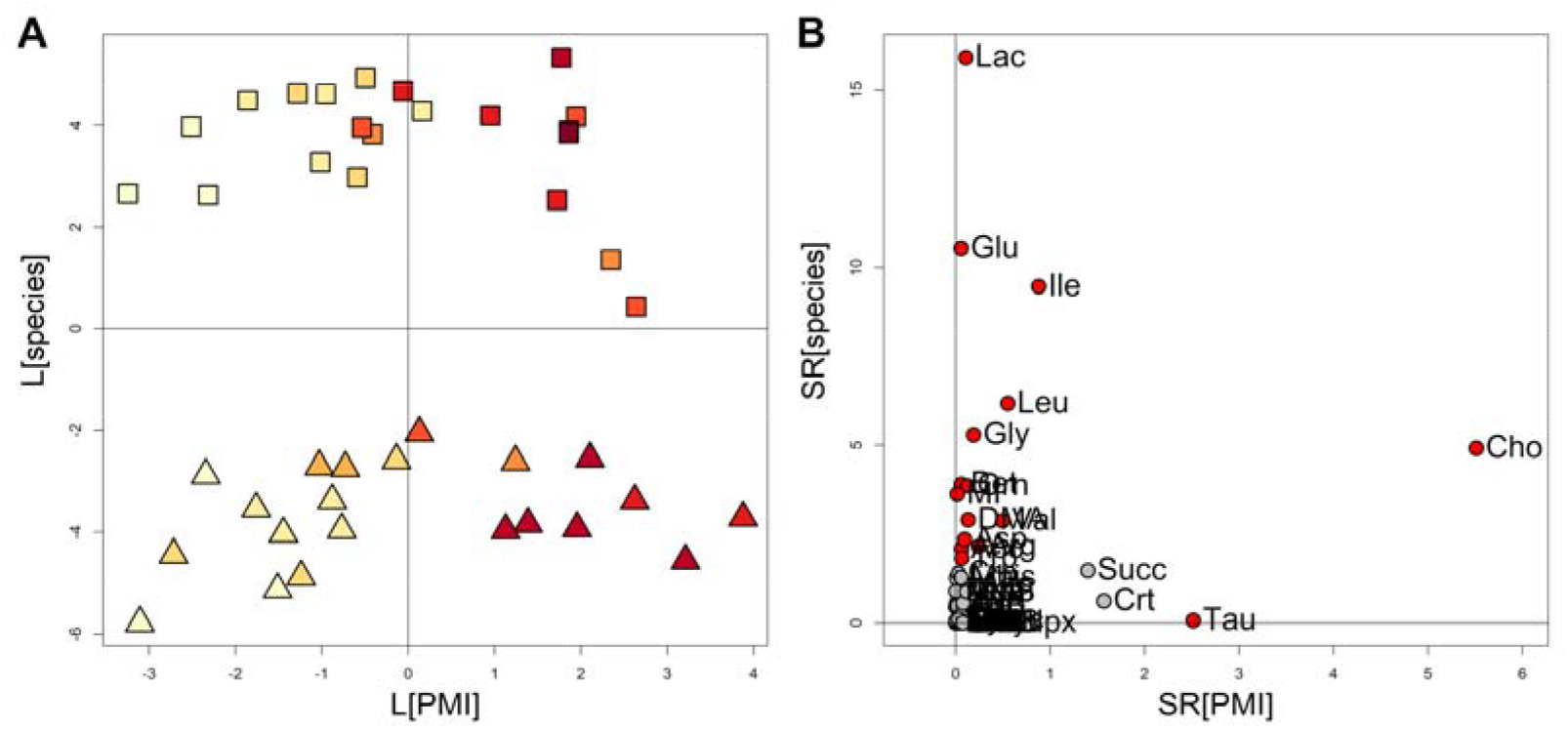
PLS for designed experiment model of the AH metabolomic profiles built to discover the effects of species and PMI. Panel A displays the PLS-scores illustrating the separation of human (boxes), and animal (triangles) samples based on species and the trends due to PMI. Panel B highlights key metabolites: species and PMI-related; significant metabolites assuming *α*=0.05 are colored in red.

To better investigate the effects of species and PMI on each single metabolite, a univariate MLR based analysis was performed. The regression coefficients of the two design factors species and PMI for the different metabolites are reported in the scatter plot of Figure 4. Interestingly, one can observe that metabolites like Lactate, Glutamate, Glycine, Betaine and Leucine play a significant role in differentiating between species, accordingly to the results of previous data analysis. On the other hand, metabolites such as Succinate, Creatine, Choline, Hypoxanthine and Taurine are more spread along the PMI-axis, with higher coefficients indicating their relevance in estimating PMI. Notably, Taurine and Hypoxanthine appear to have only a strong association with PMI independently on the factor species. Other metabolites like Serine, Phenylalanine, and others are clustered closer to the origin, indicating they have limited coefficients for both species and PMI.

**Figure 4.**
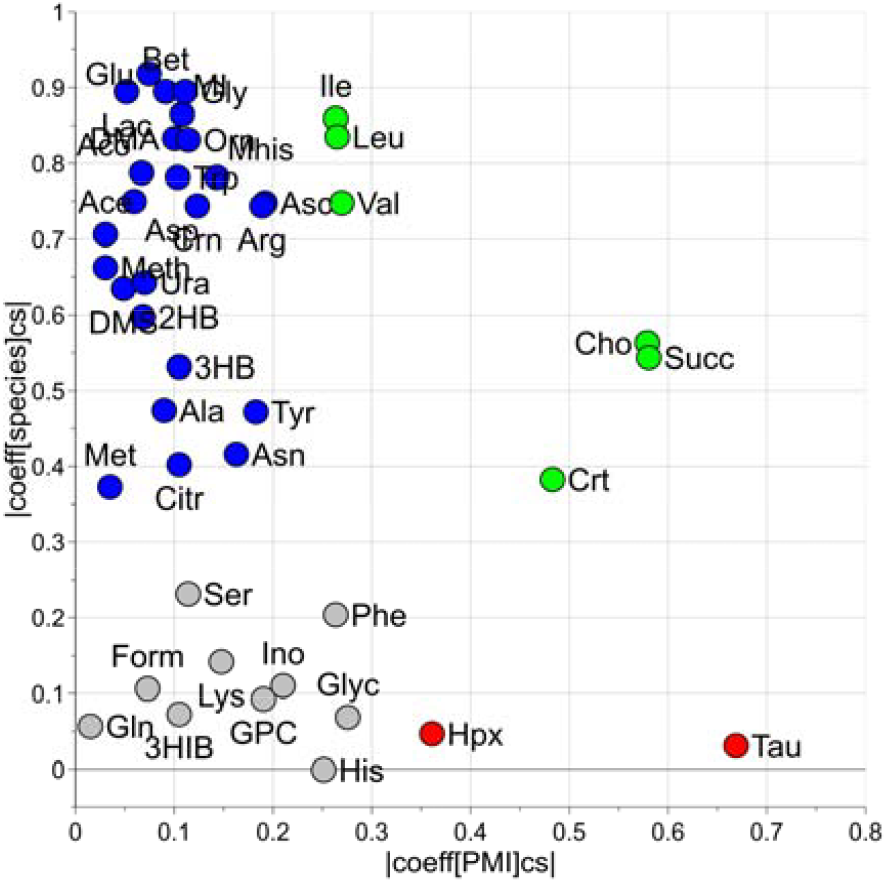
Univariate MLR based analysis: scatter plot reporting the scaled and centered coefficients of species (vertical axis) and PMI (horizontal axis) for the measured metabolites. Blue circles represent metabolites with significant coefficients only for species differentiation, green circles significant metabolites associated with both species and PMI, and red circles significant metabolites associated only with PMI.

Figure 5 illustrates boxplots of the most relevant metabolites differentiating human from animal AH, independently on their biological role in PMI. Of note, the vast majority appears to be more concentrated in the animal AH. On the contrary, concentration of a few molecules (Lactate, Creatinine, 2-Hydroxybutyrate, and Asparagine) is higher in humans AH.

**Figure 5.**
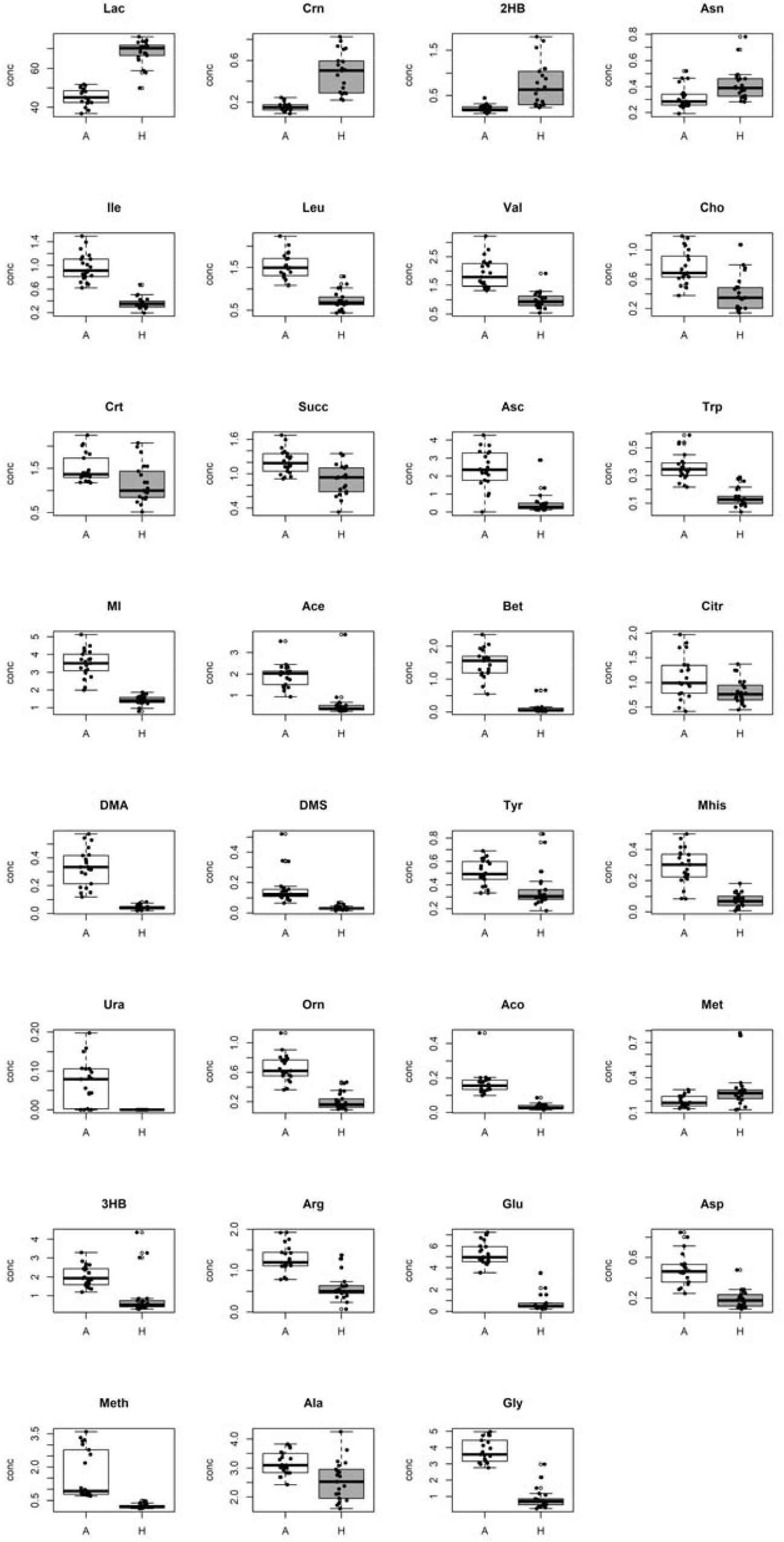
Concentration of the metabolites discovered to be significantly different between Animal (A) and Human (H) samples. Boxplots illustrate the distributions of the metabolite concentration in AH samples from the animal model (white) and human forensic cases (dark grey). In each boxplot are reported the experimental concentration (black points), the median and the interquartile range.

Interestingly, no differences in Taurine and Hypoxanthine concentrations can be seen between animal and humans as shown by Figure 6. This finding confirms that Taurine and Hypoxanthine play a role in the post-mortem independently on the species (ovine vs humans) and on the experimental setting (controlled vs real-life).

**Figure 6.**
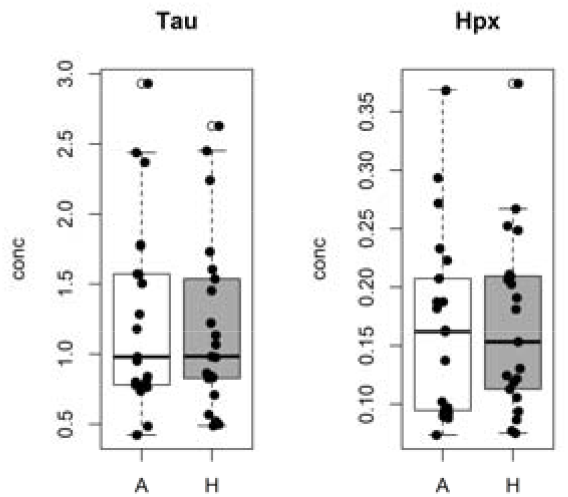
Taurine and Hypoxanthine concentrations between Animal (A) and Human (H) AH samples. Boxplots illustrate the distributions of the metabolite concentration in samples from the animal model (white) and human forensic cases (dark grey). In each boxplot are reported the experimental concentration (black points), the median and the interquartile range.

As a final step of data analysis, a PLS regression model to estimate PMI considering only the AH human samples was investigated. The model showed 2 score components, R^2^=0.886 (p=0.014), Q^2^=0.411 (p=0.002), SDEC=99 minutes and SDECV=225 minutes. Of note, SDEC and SDECV values are superposable with those calculated when human samples are projected on the animal regression model (Figure 2). The model proved that the AH metabolic content is closely related to the PMI also in the case of human samples despite the limited number of samples. Once stability selection is applied, 9 metabolites resulted to be relevant for PMI prediction. Interestingly, 6 metabolites were directly related with PMI while Glucose, Citrate, and Phenylalanine decreased over time. The plots in Figure 7 illustrate the relationships between PMI and concentration of the relevant selected metabolites.

**Figure 7.**
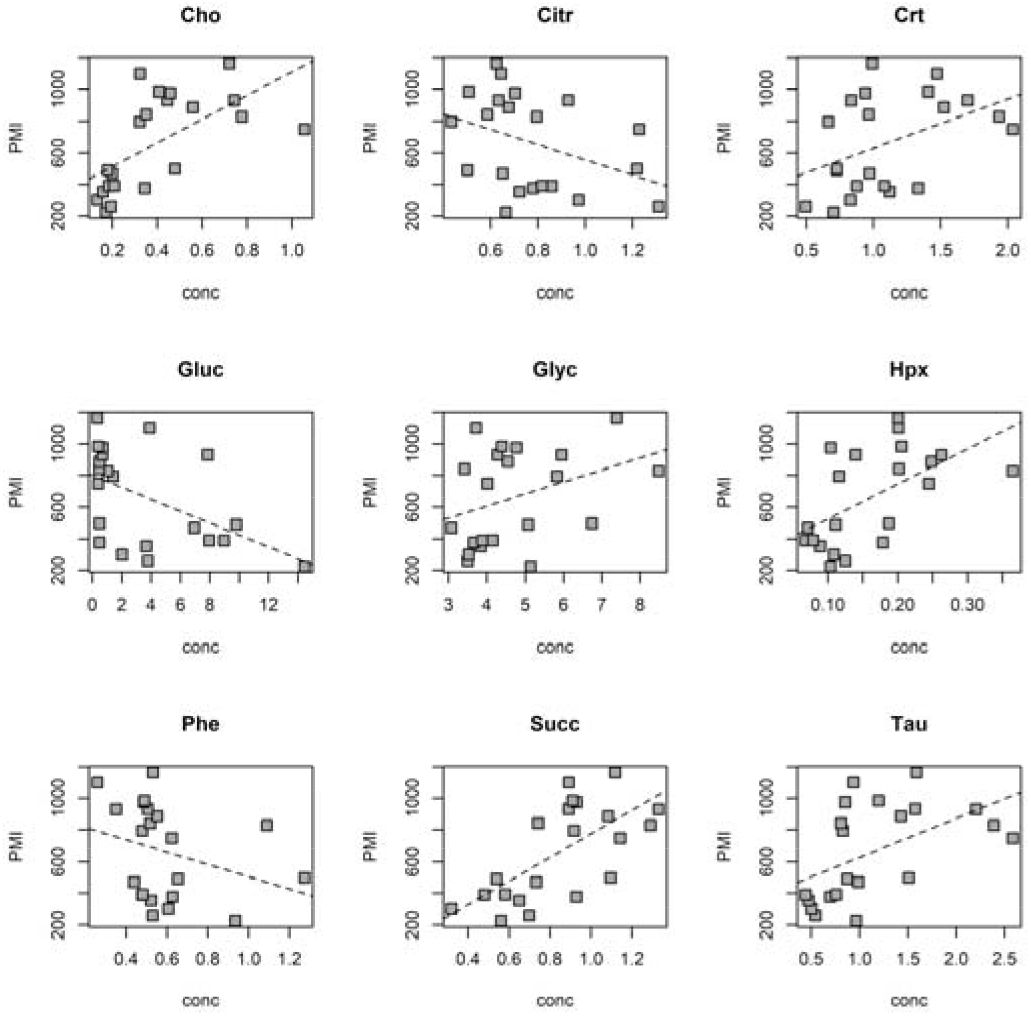
Relevant metabolites selected by PLS analysis applied to the AH human samples. The dashed line was obtained by linear regression.

## Discussion

The translation of animal model evidence into human contexts remains a fundamental challenge in scientific research, where species differences often undermine the applicability of preclinical discoveries. Reported translational success rates vary widely, ranging from 0% to 100%, often reflecting flaws in experimental design. Nevertheless, both the frequency of translational efforts and their success rates have markedly increased since 2000, as demonstrated by Leenars et al. in a recent systematic scoping review [20]. Metabolomics, a field heavily reliant on animal model experiments [21], has seen its insights translated to human applications in areas like clinical medicine [22, 23], yet this critical step remains largely unexplored in forensic research. To date, only one published paper has tried to address animal to human translation in the post-mortem metabolomics setting [24]. This study achieved only partial translation to humans, primarily due to experimental design constraints: the use of a phylogenetically distant animal model (rat) relative to humans, reliance on skeletal muscle in both species, and divergent PMI windows—up to 3 days in rats versus 3 to 19 days in humans.

Starting from our previously investigated model on *ovis aries* AH [11], our current study sought to overcome these barriers by choosing a comparable human dataset in terms of PMI. Despite the obvious differences between the animal model investigated and the human forensic setting, our results suggest a considerable similarity in the metabolomic composition of AH. Among the metabolome, three metabolites were identified only in the human samples, namely Glucose, 2-Hydroxyisovalerate, Ethanolamine. Glucose is undoubtedly linked to the carbohydrate-rich human diet also corroborated by the significantly higher lactate levels in human samples. Glucose detection in animal AH samples is indeed limited to the very early PMIs (<180 minutes) suggesting either a lower basal concentration or a fast depauperation in the post-mortem [11]. No clear explanation can be derived from these preliminary data, as the qualitative and quantitative analysis of changes in the AH metabolome was not the primary focus of this study. 2-Hydroxyisovalerate is a product of L-leucine metabolism and it has been recently identified as a potential biomarker of Residual Feed Intake (RFI) in sheep [25, 26], being RFI an efficient tool for monitoring feed efficiency. In a PLS-DA analysis comparing RFI-positive and RFI-negative cohorts, 2-Hydroxyisovalerate emerged as one of the most discriminating metabolites associated with inefficient animals.

The observed low concentrations of Glucose and 2-Hydroxyisovalerate in the aqueous humor (AH) of our samples, potentially reflecting plasma concentrations, may therefore be linked to poor breeding condition. Lastly, no mechanistic inferences can be made regarding ethanolamine, which is listed as a component of ovine plasma in the Livestock Metabolome Database (https://www.lmdb.ca/), although it has not been quantified yet.

Moreover, the post-mortem setting, which is known to represent a driving force for metabolomic modifications [10], further complicates the interspecies comparison. Despite this, our results demonstrate that the post-mortem metabolomic modifications are substantially superposable from a qualitative perspective. Worth of note the behaviour of two metabolites, namely taurine and hypoxanthine, which modifications are only influenced by the factor PMI independently on the species.

Taurine, whose antioxidant and neuroprotective role in ocular fluids is well-known [27], intriguingly resulted the best performer univariate PMI prediction in our AH animal model, resulting in a SDEP equals to 121 minutes. Hypoxanthine, a long promising metabolite in PMI estimation in VH [28] was found to be correlated with longer PMIs in the AH animal model [11].

Beside them, Succinate, Creatine, and Choline appears to be influenced by both species and PMI reflecting, at least, a partial role in the post-mortem setting. It is worth recalling that in the animal model Taurine, Hypoxanthine, Succinate, Creatine, and Choline showed a continuous and monotonous trend over the entire time-window investigated [11]. In this work, the same metabolites are influenced by PMI despite the behaviour of some of them is affected by the factor species. Focusing on the forensic insights of this work, despite the low number of samples, a multivariate regression model based only on human samples was built displaying similar errors in PMI prediction when compared to the more robust animal vs human model. Of note, 9 metabolites resulted to be relevant for PMI prediction in the human model, including the above-mentioned 5 metabolites plus Glucose, Glycine, Citrate, and Phenylalanine. Three out of nine relevant metabolites showed negative correlation whereas the other six accumulated over time indicating a dynamic activity of the AH metabolism in the post-mortem.

This work represents a fundamental step in the translational research. Animal models are indeed widely used due to practical and/or ethical limitations but are rarely translated into human samples, especially in the post-mortem setting. One of the general criticisms about animal models is the aprioristic assumption that the baseline differences hamper the human translation. In our work we were able to demonstrate that part of the metabolome is indeed severely affected by the species although only from the quantitative perspective. At the same time, another significant portion of the metabolome was found to be linked to the independent variable under investigation culminating in two important metabolites indifferently from the factor species. Another point to be stressed is that the investigated animal model although certainly complex is relatively far from humans’ biological reality highlighting that post-mortem related phenomena are driven by transversal factors (i.e., environmental temperature, metabolism backbone under hypoxic conditions and microbial co-metabolism) resulting in shared metabolomic modifications among different models.

This paper, which represents the ultimate work of a research line on post-mortem metabolomics on ocular matrices [11-13], was designed with ^1^H NMR which is known to be a robust and reproducible tool. An interesting follow up study could be the use of mass spectrometry in order to test the approach on an analytical platform which is inherently less robust but much more sensitive to finally corroborate these results.

Our study presents a number of limitations. Firstly, human dataset was based on hospital deaths resulting in an elder population. This was due to need of very early PMI to match our previous AH animal model. The human forensic dataset was then based on a population with several comorbidities, enhancing the difference with the highly controlled animal model (young and healthy sheep).

Secondly, the sample size was limited to 21 samples collected from 11 individuals, limiting the inter-individual variability in the human population. AH from eyes of the same individuals were collected at different time-points allowing PMI-related modifications to occur. Lastly, AH represents a biofluid of limited practical interest in the forensic scenario due to its fast dehydration. In the broader picture of PMI investigation, it was chosen because it represents the ocular biological matrix where the post-mortem modifications begin in the eye compartment and, reasonably, the base for other ocular matrices changes.

The same translational approach is currently under study by our research group on VH. The importance of translating animal model results [VH] on human VH is linked to the possibility of investigating a wider time-window (up to 100 hours after death) of more significant practical forensic interest.

## Declarations

### Authors’ contributions

### Data availability

The datasets generated during and/or analysed during the current study are available from the corresponding author on reasonable request.

### Ethics declarations

As samples were obtained during Judicial Autopsies informed consent was obtained through local Prosecutor Office, office, which acted in what was believed to be the closest interest of the deceased being the samples completely anonymised.

